# Pharmacological Depletion of Retinal Mononuclear Phagocytes is Neuroprotective in a Mouse Model of Mitochondrial Optic Neuropathy

**DOI:** 10.1101/2025.08.11.669571

**Authors:** Avital L. Okrent Smolar, Rahul Viswanath, Howard M. Bomze, Ying Hao, Sidney M. Gospe

## Abstract

**Purpose:** The *Vglut2-Cre;ndufs4*^*loxP/loxP*^ mouse strain with retinal ganglion cell (RGC)-specific mitochondrial complex I dysfunction develops severe RGC degeneration by postnatal day 90 (P90), with accompanying retinal mononuclear phagocyte (MNP) accumulation. We have reported that continuous exposure to hypoxia partially rescues RGC death in these mice, with minimal effect on MNP abundance. We hypothesized that pharmacological depletion of MNPs with the colony-stimulating factor-1 receptor inhibitor pexidartinib would enhance RGC neuroprotection by hypoxia.

**Methods:** Iba1^+^ retinal MNP depletion was assessed in C57Bl/6J mice fed control or pexidartinib-infused chow beginning at P25. Subsequently, *Vglut2-Cre;ndufs4*^*loxP/loxP*^ mice and control littermates were raised under normoxia or hypoxia and fed control or pexidartinib chow from P25 to P90. The neuroprotective effect of pexidartinib and hypoxia alone and in combination was assessed by quantifying RGC soma and axon survival in retinal flat mounts and optic nerve cross sections.

**Results:** Pexidartinib completely depleted retinal MNPs within one week of treatment. Untreated *Vglut2-Cre;ndufs4*^*loxP/loxP*^ mice exhibited the expected ∼50% reduction of RGC soma and axon survival at P90 (p<0.0001 for both). Hypoxia or pexidartinib monotherapy each reduced RGC degeneration by more than one-half, while their combination resulted in complete RGC neuroprotection (p<0.001 for all three treatments). Normal myelination patterns were restored in mice receiving dual therapy.

**Conclusions:** Pexidartinib effectively depletes retinal MNPs and is neuroprotective in the setting of severe RGC mitochondrial dysfunction. This therapeutic effect is additive to that of hypoxia. Combating retinal neuro-inflammation may therefore be a useful adjunct therapy in mitochondrial optic neuropathies like Leber hereditary optic neuropathy.

## INTRODUCTION

Leber hereditary optic neuropathy (LHON) is the most common heritable disease resulting from primary mutations in the mitochondrial DNA (mtDNA).^1^ It is characterized by profound bilateral vision loss in most patients due to rapidly progressive degeneration of retinal ganglion cells (RGCs), which are particularly sensitive to the mitochondrial complex I dysfunction produced by LHON-associated mutations.^2^ Effective treatment for LHON and other mitochondrial optic neuropathies is an important unmet need. Although there is evidence in the literature of some clinical benefit from treatment with oral idebenone to improve mitochondrial function, most LHON patients continue to have severe visual impairment despite treatment.^3-5^ Furthermore, while early gene therapy trials for LHON have shown some promise,^6,7^ a genetically heterogenous disease like LHON would require the development of specific gene therapies for each mutated complex I subunit. The identification of an effective mutation-independent treatment strategy would be a significant advancement for patient care.

In an effort to develop an optimal preclinical mouse model of rapidly progressive mitochondrial optic neuropathy, we have generated a genetically modified mouse line in which severe complex I deficiency is induced within RGCs in a cell-specific manner.^8^ This model utilizes Cre recombinase driven by the Vglut2 promoter to delete *floxed* alleles of the nuclear-encoded complex I accessory subunit *ndufs4* within RGCs. Deletion of this gene, which is mutated in some forms of Leigh syndrome,^9^ destabilizes complex I and decreases its enzymatic activity by >50% in the retina and other tissues.^10-12^ This severe compromise of complex I function results in rapid degeneration of RGCs that begins around postnatal day 45 (P45) and becomes profound by P90.^8^ Additionally, as an indicator of retinal neuro-inflammation, the mice exhibit inner retinal accumulation of Iba1^+^ mononuclear phagocytes (MNPs), representing proliferation of native microglia and/or infiltration of macrophages derived from peripheral monocytes. The onset of acute RGC loss just after the mice reach sexual maturity is akin to many human cases of LHON and supports the use of this mouse line as a preclinical model for the disease.

We have previously reported that when the *Vglut2-Cre;ndufs4*^*loxP/loxP*^ mouse mice are raised under continuous hypoxia (11% O_2_) beginning at P25, they exhibit marked rescue of RGC axons and somas, with a neuroprotective effect persisting to at least P90.^13^ However, despite reducing RGC degeneration, hypoxia has minimal, if any, effect on the retinal accumulation of MNPs.^13^ Interestingly, it has recently been reported that pharmacological depletion of MNPs results in a significant prolongation of lifespan and inhibition of brain lesion formation in a Leigh syndrome mouse model with global deletion of *ndufs4*.^14,15^ Furthermore, retinal MNPs have been directly implicated in the pathogenesis of RGC loss in animal models of another optic neuropathy, glaucoma.^16,17^ Therefore, we were interested to investigate whether depletion of MNPs could reduce RGC degeneration in our mitochondrial optic neuropathy model. Herein we report that systemic administration of the colony-stimulating factor-1 receptor (CSF-1R) inhibitor pexidartinib (PLX3397) causes rapid and profound elimination of MNPs from mouse retinas and has a neuroprotective effect on *ndufs4*-deficient RGCs that is equivalent to that of continuous hypoxia. Most notably, the salutary effect of pexidartinib was additive to that of hypoxia, with a 100% rescue of RGC somas and axons observed in mice receiving both therapies.

## METHODS

### Animals

All animal experiments adhered to the ARVO Statement for the Use of Animals in Ophthalmic and Vision Research, following a protocol approved by the Institutional Animal Care and Use Committee of Duke University. Wild type C57Bl/6J mice were purchased from Charles River Laboratories (Wilmington, MA). *Vglut2-Cre;ndufs4*^*loxP/loxP*^ mice and control littermates were generated as previously described^8^ and maintained on a C57Bl/6J background.

Starting at P25, shortly after weaning, the mice were given *ad libitum* access to D11112201 OpenStandard diet (Research Diets, Inc., New Brunswick, NJ) with or without 400 mg/kg of pexidartinib (PLX3397, MedKoo Biosciences, Durham, NC). This was the sole food source of the mice until euthanasia for collection of ocular tissues.

### Continuous Hypoxia

Hypoxia treatment was performed as previously described.^13^ Briefly, mouse cages assigned to hypoxia groups were kept within a hypoxia chamber (A-Chamber animal cage enclosure; BioSpherix, Ltd., Parish, NY) with ambient PO_2_ reduced to 11% by pumping in nitrogen gas to displace the oxygen. The mice were maintained in the hypoxia chamber until P90, under a 12-hour light/dark cycle. Cages with cohorts maintained under normoxia remained in their standard racks in the same animal facility.

### Antibodies

The following antibodies were used for immunofluorescence experiments: rabbit polyclonal anti-RBPMS1 (1:1000, NBP2-20112; Novus Biologicals, Englewood, CO), rabbit polyclonal anti-Iba1 (1:1000, 019–19741; Fujifilm Wako Chemicals Corporation, Richmond, VA), and mouse monoclonal anti-Tuj1 (1:5000; MAB11905; Thermo Fisher Scientific, Waltham, MA). Secondary antibodies against the appropriate species conjugated to Invitrogen Alexa Fluor 488, 568, or 647 (1:5000 dilution) were purchased from Thermo Fisher Scientific. Cell nuclei were stained using Hoechst 33342 (5 µg/ml; Thermo Fisher Scientific).

### Histological Techniques

Immunofluorescence analyses were performed as previously described.^8^ Briefly, enucleated eyes obtained from euthanized mice were fixed overnight in 4% paraformaldehyde. Flat mounts were made from isolated retinas by making four radial cuts from the edge to the equator of each retina and then blocked in 5% donkey serum in PBS with 0.3% Triton X-100. The retinas from C57Bl/6J mice were co-incubated with anti-Iba1 and anti-Tuj1 primary antibodies. For the RGC rescue experiment, one quadrant from each retina was removed and incubated with anti-Iba1 and anti-Tuj1 primary antibodies, while the remaining three quadrants were incubated in anti-RBPMS1 primary antibody. After 5 days of incubation at 4°C, the retinas were washed and incubated with secondary antibodies in block overnight at 4°C. The retinas were then washed, mounted on glass slides with the RGC layer facing up in Fluoromount-G (SouthernBiotech, Birmingham, AL) and coverslipped.

Images were acquired on a confocal microscope (Eclipse 90i and A1 confocal scanner; Nikon, Tokyo, Japan) with a 60× objective (1.4 NA Plan Apochromat VC; Nikon) using Nikon NIS-Elements software. Images 45,000 µm^2^ in area were obtained in each quadrant at three distances from the optic nerve head: 0.5, 1.0, and 1.5 mm. For RGC soma quantification, images were obtained as z stacks spanning the ganglion cell layer. For MNP quantification, z stacks were obtained from the base of the inner nuclear layer to the inner (vitreous) surface of the retina. RBPMS1^+^ RGC somas and Iba1^+^ MNPs were manually counted using ImageJ software (National Institutes of Health, Bethesda, MD). The abundance of each cell type was averaged among all quadrants at each distance from the optic nerve head.

To assess RGC axons, optic nerves were obtained from the same mice and fixed in 2% paraformaldehyde and 2% glutaraldehyde in PBS for 2 hours at room temperature. Samples were embedded in EMbed 812 resin mixture (Electron Microscopy Sciences, Hatfield, PA) and sectioned on an ultramicrotome (LKB Ultratome V; Leica, Wetzlar, Germany) using a glass knife. Cross-sections of 0.27-µm thickness were stained with 1% methylene blue. For each optic nerve specimen three cross-sections were imaged using a Nikon ECLIPSE Ti2 microscope and NIS-Elements imaging software, with sufficient images obtained using a 60× (oil) objective to cover the entire area of the nerve cross section. The images were stitched into a composite image representing the entire cross section. Axon count analysis was performed using the AxoNet plugin for ImageJ.^18^ The final axon count was divided by the total optic nerve area to determine the mean axon density and then averaged over the three cross-sections for each nerve.

For ultrastructural analysis, the same optic nerve specimens were thinly sectioned (60–80 nm), collected on copper grids, counterstained with uranyl acetate and Sato’s lead, and then examined using a transmission electron microscope (JEM-1400; JEOL USA, Peabody, MA) at 60 kV. Images were collected using an Orius charge-coupled device camera (Gatan, Pleasanton, CA).

### Experimental Design and Statistical Analysis

Histological assessment of MNP depletion from the retinas of male and female wild type C57Bl/6J mice was performed on 6-8 retinas from each group at each time point. Quantification of RGC soma and axon survival and of MNP abundance was performed on *Vglut2-Cre;ndufs4*^*loxP/loxP*^ mice and *Vglut2-Cre;ndufs4*^*loxP/+*^ littermate controls, with both sexes represented in each group. In these experiments, 10 to 16 retinas or optic nerves were analyzed for each combination of genotype, O_2_ concentration, and chow. While performing cell counts, the observer was masked to the identity of each sample. Statistical comparisons between groups were made with the Wilcoxon rank-sum test to account for non-parametric data. All data analysis for this study was performed with Prism 10 software (GraphPad, San Diego, CA). Data are presented graphically as mean ±LSEM, and biological replicates are depicted as individual data points on the bar graphs.

## RESULTS

Based on two recent reports that treatment of the *ndufs4*^*-/-*^ mouse model of Leigh syndrome with pexidartinib resulted in a several-fold prolongation of the lifespan of the animals, we were interested to determine whether the drug would prove neuroprotective in our mouse strain with RGC-specific deletion of the gene. Pexidartinib is a tyrosine kinase inhibitor that targets CSF-1R to deprive MNPs of a key survival signal. It has been shown to cross the blood-brain barrier, effectively depleting MNPs from brain as well as retinal tissue.^19,20^ However, there have been conflicting data regarding whether the efficacy of pexidartinib in depleting Iba1^+^ MNPs from the mouse brain is sex-dependent, with some reports describing a much more pronounced effect in male mice compared to females.^21,22^ In order to determine how rapidly and completely pexidartinib eliminates MNPs from the retina, and whether this would be impacted by the sex of the mice, we first administered the drug to young wild type C57Bl/6J mice beginning just after weaning at P25 via drug-infused chow (400 mg/kg).^22^ Retinal tissue was harvested after either 1 week (P32) or one additional month (P60) to determine the extent and durability of MNP depletion. We observed near-complete loss of Iba1^+^ cells from both the inner and outer retina after just one week of treatment (Figure 1A). Because only those MNPs in the vicinity of RGCs would be likely to impact their survival in a mitochondrial optic neuropathy, we limited our quantitative analysis of MNP abundance to those localizing to the inner retina (from the junction of the inner nuclear layer and inner plexiform layer to the internal limiting membrane). With rare exception, we observed complete loss of inner retinal Iba1^+^ cells at P32, with no difference between male and female mice (Figure 1B). Continued administration of pexidartinib achieved a durable elimination of MNPs, with none observed in any of the retinas at P60, irrespective of sex (Figure 1C).

**Figure 1.**
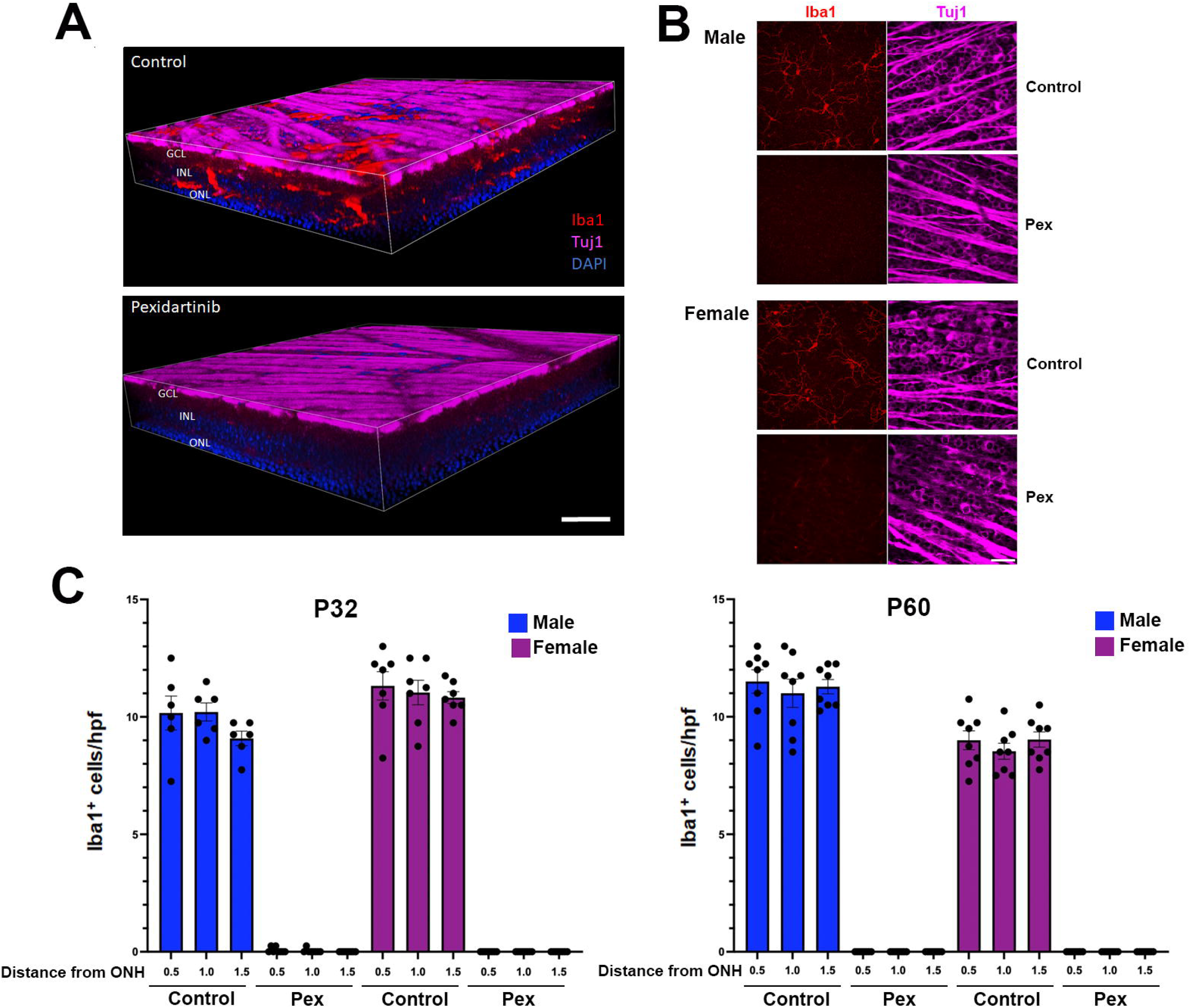
Systemic administration of pexidartinib efficiently depletes retinal mononuclear phagocytes from wild type mice. (A) Representative 3-dimensional reconstructions of confocal microscopy z-stacks of retinal flat mounts from wild type C57Bl/6J mice administered control (top) or pexidartinib-infused chow (bottom) for 1 week. Immunolabeling for β3-tubulin (Tuj1; magenta) highlights the retinal nerve fiber layer comprised by RGC axons, while mononuclear phagocytes (MNPs) are immunolabeled in red for Iba1. The nuclear layers of the retina are labeled with DAPI (blue). Pexidartinib treatment eliminated Iba1^+^ MNPs from both the inner retina (those residing between the inner nuclear layer and ganglion cell layer) and the outer retina (those deep to the inner nuclear layer). GCL, ganglion cell layer; INL, inner nuclear layer; ONL, outer nuclear layer. Bar, 40 µm. (B) Representative flat mounts of P32 retinas of wild type male and female mice, after one week of treatment with control or pexidartinib chow. MNPs are labeled with Iba1 (red) and the nerve fiber layer with Tuj1 (magenta). (C) Quantification of the abundance of Iba1^+^ MNPs per high power field (hpf) in the inner retina at distances of 0.5, 1.0, and 1.5 mm from the optic nerve head, demonstrating near-complete depletion in mice of both sexes treated with pexidartinib. Bar, 40 µm. Error bars depict mean ± SEM. (D) Comparison of Iba1^+^ MNP abundance in wild type mice of both sexes at P60 that have been treated for an additional month with control or pexidartinib chow.

Because RGC loss in the *Vglut2-Cre;ndufs4*^*loxP/loxP*^ mouse model of mitochondrial optic neuropathy does not commence until after P30 and progresses similarly in mice of both sexes,^7^ we proceeded to test the effect of pexidartinib treatment beginning at P25, using both male and female littermates. The mutant mice live longer than global *ndufs4* knockout mice, but do not survive much beyond P90 due to progressive neurological dysfunction arising from complex I deficiency in those glutamatergic neurons in the brain expressing Vglut2-driven Cre recombinase.^8,23^ We therefore selected P90 as the terminal timepoint at which ocular tissues would be harvested. Pexidartinib-infused chow or control chow were administered as the sole sources of food to the experimental *Vglut2-Cre;ndufs4*^*loxP/loxP*^ mice and control *Vglut2-Cre;ndufs4*^*loxP/+*^ littermates that are heterozygous for loss of *ndufs4* and previously shown to be aphenotypic.^13^ In order to compare any neuroprotective effect of pexidartinib to that of hypoxia and to determine any added benefit when the two treatments were combined, half of the mice from each group were raised under normoxia (21% O_2_) or continuous hypoxia (11% O_2_) beginning at P25. Whereas the *Vglut2-Cre;ndufs4*^*loxP/loxP*^ mice fed a control diet and raised under normoxia demonstrated the expected severe stiffened posture and ataxia by P90, those treated with pexidartinib and/or hypoxia exhibited a body posture and activity level grossly indistinguishable from their heterozygous littermates, consistent with the neurological improvement reported in studies of global *ndufs4*^*-/-*^ mice administered either treatment.^15,24^ All mice treated with pexidartinib exhibited an acquired depigmentation of their hair, previously observed in mice and a known side effect of the drug in humans^15,25^ (Figure 2A).

**Figure 2.**
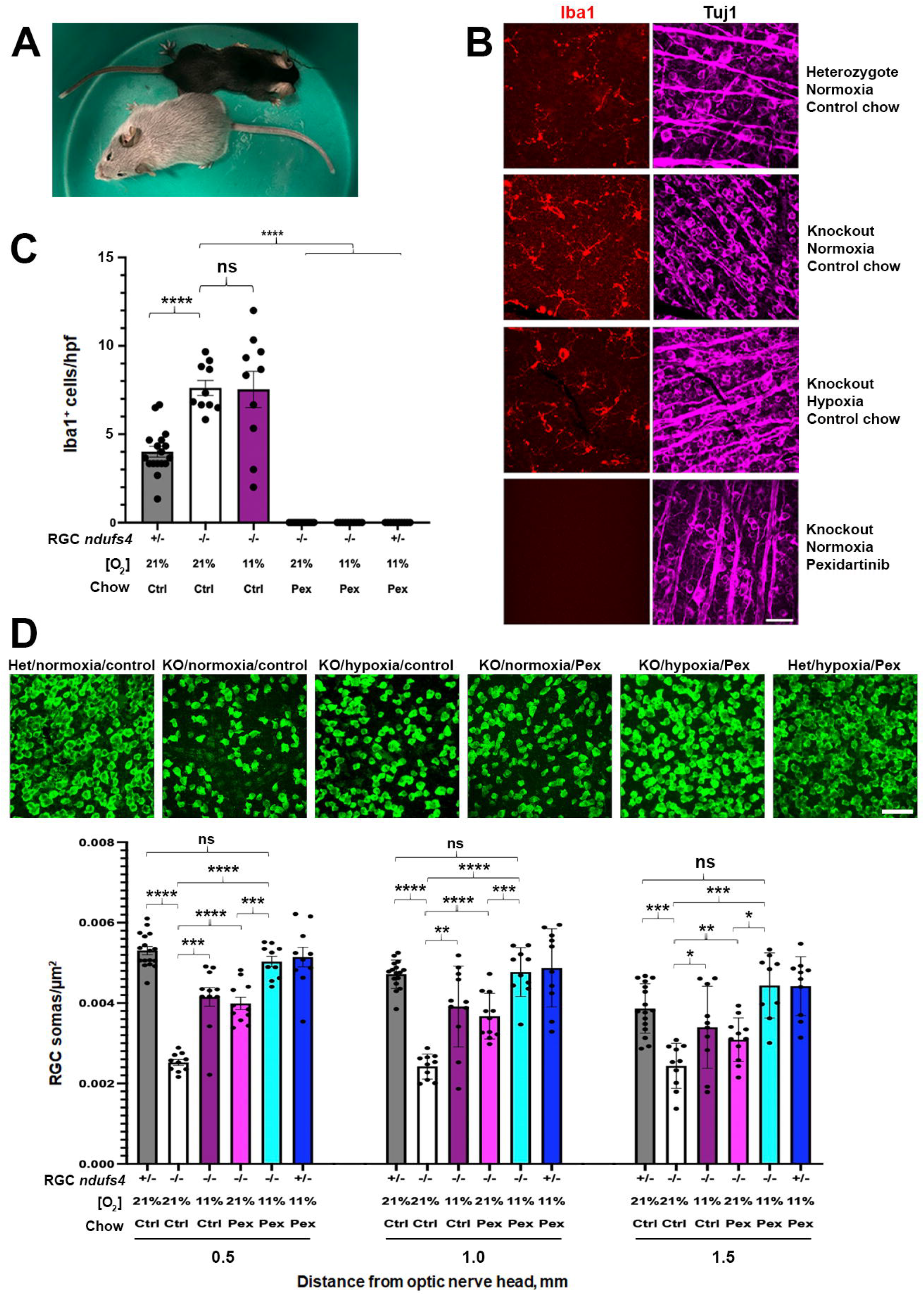
Pexidartinib treatment of *Vglut2-Cre;ndufs4*^*loxP/loxP*^ mice produces a neuroprotective effect on retinal ganglion cell soma degeneration that is additive to that of hypoxia. (A) Photograph of *Vglut2-Cre;ndufs4*^*loxP/loxP*^ littermates at P90. The top mouse received control chow and has become weak, with splayed limbs. The bottom mouse received pexidartinib chow and demonstrates a normal posture. Graying of the coat is a side effect of high dose pexidartinib. (B) Representative images of Iba1-labeled (red) retinal flat mounts from a heterozygous *Vglut2-Cre;ndufs4*^*loxP/loxP*^ control mouse and from *Vglut2-Cre;ndufs4*^*loxP/loxP*^ mice that are untreated or received hypoxia or pexidartinib treatment. Tuj1 (magenta) was used to confirm image capture at the level of RGCs. Bar 40 µm. (C) Comparison of Iba1^+^ MNP abundance between experimental cohorts. *ndufs4*^*+/-*^ indicates deletion of one allele of *ndufs4* from RGCs (*Vglut2-Cre;ndufs4*^*loxP/+*^ mice), and *ndufs4*^*-/-*^ indicates deletion of both alleles (*Vglut2-Cre;ndufs4*^*loxP/loxP*^ mice). There is an accumulation of Iba1^+^ MNPs in the inner retina in untreated mice with *ndufs4*-deficient RGCs that is unaffected by hypoxia treatment. Complete depletion of MNPs from the inner retina was observed in all cohorts receiving pexidartinib. (D) Representative images of RBPMS1-labeled RGC somas (green) obtained at a distance of 1.0 mm from the optic nerve head in retinal flat mounts from each experimental cohort at P90. The graph below demonstrates a significant reduction of RGC soma loss in *Vglut2-Cre;ndufs4*^*loxP/loxP*^ mice treated with hypoxia or pexidartinib; the combined treatment restores RGC survival to the level observed in heterozygous *Vglut2-Cre;ndufs4*^*loxP/loxP*^ control mice that were untreated or received the dual therapy. Graphs depict mean ± SEM. Biological replicates are indicated as individual data points. *, p≤0.05; **, p≤0.01; ***, p≤0.001; ****, p≤0.0001; ns, not significant.

The abundance of Iba1^+^ MNPs in the inner retina was assessed in retinal flat mounts from each group (Figure 2B). Similar to our prior observations from retinal cross sections, MNPs were observed at a frequency 2-fold higher in untreated *Vglut2-Cre;ndufs4*^*loxP/loxP*^ mice at P90 compared to heterozygous controls, and this increase was not mitigated by hypoxia treatment (Figure 2C). Pexidartinib treatment resulted in a complete elimination of inner retinal MNPs at P90 in all groups receiving the drug.

Retinal flat mounts were immunolabeled with the RGC marker RBPMS1 to allow quantification of RGC soma survival in each group (Figure 2D). Consistent with our prior observations, the untreated *Vglut2-Cre;ndufs4*^*loxP/loxP*^ mice exhibited approximately 50% loss of RGC somas at P90 compared to heterozygous littermates. Also as noted previously, continuous hypoxia prevented more than half of this attrition in *Vglut2-Cre;ndufs4*^*loxP/loxP*^ mice. Pexidartinib treatment resulted in a similarly robust but incomplete rescue of RGCs. Most remarkably, the dual treatment with hypoxia and pexidartinib showed an additive neuroprotective effect, fully restoring RGC soma density to that seen in the heterozygous controls. Notably, heterozygous mice administered the dual treatment also exhibited normal RGC survival, indicating that neither treatment was deleterious to healthy RGCs without mitochondrial dysfunction.

We then assessed the optic nerves of these mice to assess the neuroprotective effect of each treatment on RGC axons (Figure 3A). Similar to RGC somas, there was a ∼50% reduction in axon density in untreated *Vglut2-Cre;ndufs4*^*loxP/loxP*^ mice at P90. A partial but significant increase in RGC axon survival was observed in the knockout mice receiving monotherapy with either hypoxia or pexidartinib, while axon survival was restored to normal levels in those mice receiving dual therapy. Optic nerve ultrastructure was then assessed by transmission electron microscopy (Figure 3B). Characteristic myelination abnormalities were observed in specimens from the untreated *Vglut2-Cre;ndufs4*^*loxP/loxP*^ mice, including thickening or duplication of myelin sheaths or incomplete enclosure of axons by myelin. Dual therapy with hypoxia and pexidartinib prevented the development of pathological myelination patterns, while partial improvement was observed when the experimental mice were administered either treatment in isolation.

**Figure 3.**
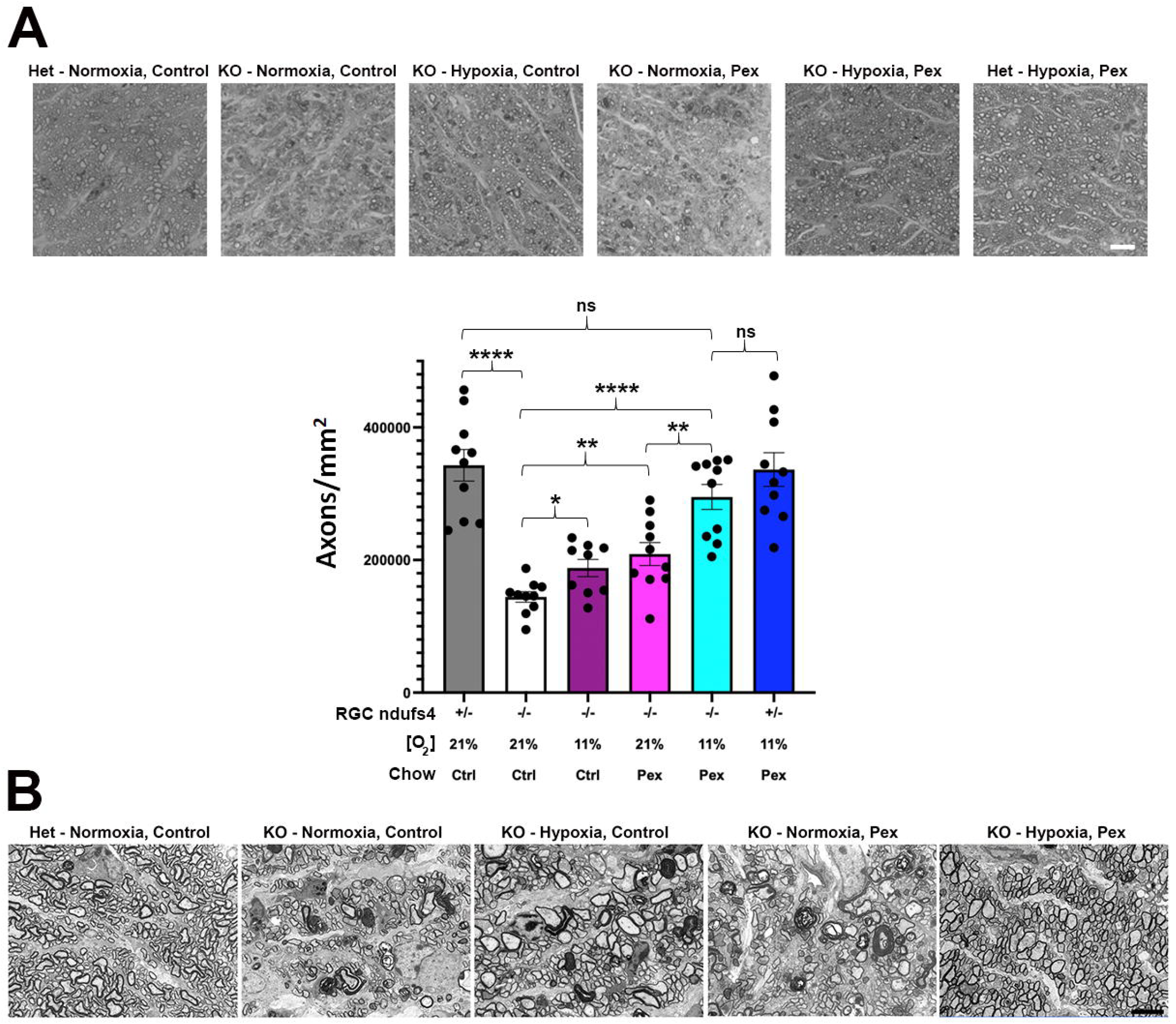
Hypoxia and pexidartinib reduce RGC axonal degeneration in *Vglut2-Cre;ndufs4*^*loxP/loxP*^ mice in an additive manner. (A) Representative light micrographs of optic nerve cross sections stained with methylene blue. The genotype within RGCs [heterozygous (Het) or knockout (KO) for *nduf4*] and treatment of each cohort is indicated above each panel. Bar, 10 µm.The graph below shows RGC axon densities in optic nerve cross sections determined using AxoNet automated axon quantification. *ndufs4*^*+/-*^ indicates deletion of one allele of *ndufs4* (*Vglut2-Cre;ndufs4*^*loxP/+*^ mice), and *ndufs4*^*-/-*^ indicates deletion of both alleles within RGCs (*Vglut2-Cre;ndufs4*^*loxP/loxP*^ mice). The O_2_ concentration is depicted for each group, as is the type of chow administered. Individual data points depict biological replicates. Data are presented as mean ± SEM. Statistical comparisons between groups are indicated above the bars, with the following significance designations: *, p≤0.05; **, p≤0.01; ***, p≤0.001; ****, p≤0.0001; ns, not significant. (C) Electron micrographs (5000X) of optic nerve cross sections at P90 demonstrate preservation of normal axon morphology and abundance in *Vglut2-Cre;ndufs4*^*loxP/loxP*^ mice treated with both hypoxia and pexidartinib, while those receiving monotherapy demonstrate a partial reduction of abnormal myelination compared to the untreated *Vglut2-Cre;ndufs4*^*loxP/loxP*^ mice. Bar, 5 µm.

## DISCUSSION

In this study, we observed that mice undergoing systemic treatment with pexidartinib exhibited a rapid and virtually complete depletion of MNPs within 1 week of starting the therapy. This depletion persisted over two months of chronic treatment and was not dependent on the sex of the mice. The drug itself and the consequent loss of retinal MNPs did not adversely affect the survival of RGCs. When applied to mice with RGC-specific complex I dysfunction, pexidartinib produced a significant, though partial, rescue of RGC soma and axon degeneration, similar in magnitude to that observed when the mice were instead treated with continuous hypoxia (11% O_2_). Most remarkably, the neuroprotective effects of each of these treatments proved to be additive, with the combined therapy resulting in essentially complete rescue of RGCs at P90, suggesting that the treatments work through independent mechanisms.

The most intuitive approach to treating a mitochondrial disease is to mitigate the effects of mitochondrial dysfunction directly. This has been the rationale behind LHON treatments such as idebenone and gene therapy.^26^ Idebenone, a synthetic Coenzyme Q10 analog, is thought to promote RGC neuroprotection by delivering electrons to complex III, thereby bypassing the dysfunctional complex I to augment mitochondrial ATP production while also serving as a potent antioxidant to reduce oxidative stress.^27^ It is commonly administered to LHON patients within the first several years of vision loss, and studies have suggested a modest benefit in visual outcome when compared to placebo or natural history, possibly dependent on specific mtDNA mutations.^4,5^ Gene therapy trials have aimed to restore normal complex I function in RGC mitochondria by using adeno-associated virus to deliver wild type versions of mutated mitochondrial genes, thus far limited to two genes, *mtND4* and *mtND1*.^7,28^ Similar to idebenone, the gene therapies have shown generally positive outcomes, although most subjects have remained legally blind.^6^ Hypoxia, which we have previously described as protective against degeneration of RGC somas and axons in our mouse model of RGC *ndufs4* deficiency,^13^ is also likely to have a direct effect on RGCs. While the neuroprotective mechanism of hypoxia remains under investigation, it likely occurs via reducing the production of reactive oxygen species and/or promoting the forward transfer of electrons within affected mitochondria.^29,30^ Nevertheless, despite its effectiveness at slowing RGC degeneration, hypoxia has shown no more than minimal impact in reducing accumulation of Iba1^+^ MNPs, at least in the retina.^13^

In contrast to environmental hypoxia, the neuroprotective effect of pexidartinib is likely to be indirect, via its actions on MNPs. In addition to our own observations of retinal MNP accumulation in the RGC-specific *ndufs4* deletion model, prior analysis of the retinas of mice with global deletion of *ndufs4* revealed a substantial increase in the expression of cytokines and other genes related to innate immunity, even prior to the onset of RGC loss.^31^ It should be noted that in our RGC-specific disease model, the MNPs themselves remain metabolically intact, so their accumulation in the retina is presumably secondary to signaling from stressed RGCs. The role that MNPs, particularly microglia, play in the pathophysiology of degenerative retinal diseases is controversial, with both adaptive and pathological mechanisms having been described.^32,33^ In some contexts, such as retinitis pigmentosa, MNPs may be critical to clearing debris and secreting pro-survival signals, whereas in other diseases, MNPs may exhibit neurodegenerative molecular signatures and promote damage to retinal neurons, as recently described in a mouse glaucoma model.^17,34^ Our observation of improved survival of RGCs with mitochondrial impairment when MNPs are eliminated from the retina would suggest that, on balance, MNPs exacerbate the pathogenesis of mitochondrial optic neuropathy. This is consistent with a number of recent publications demonstrating a positive effect of MNP depletion on various CNS pathologies such as traumatic brain injury and other neurodegenerative diseases.^21,35-37^

As an inhibitor of the CSF-1R, pexidartinib deprives immune cells of a critical survival signal, resulting in depletion not only of native microglia but also myeloid and even a subset of lymphoid cells by suppressing progenitor cell lines in the spleen, bone marrow, and blood.^38^ It is therefore uncertain whether the therapeutic effect of pexidartinib on RGCs in the *ndufs4* mouse model is mediated by depletion of retinal microglia or a prevention of infiltration of circulating immune cells. Defining the relevant MNP population that accumulates in the retinas of mice with RGC-specific complex I deficiency will be an important future direction that could ultimately lead to more targeted therapies. Because overlapping immunophenotypes (including Iba1 positivity) make it challenging to distinguish microglia from infiltrating macrophages using conventional immunohistochemistry methods, lineage tracing experiments with genetically-modified mouse lines may be required for a definitive determination.^39^ Potentially instructive is a recent report that the *Csf1r*ΔFIRE mutation that eliminates microglia but not monocyte-derived MNPs did not reduce the formation of brain lesions and neurologic dysfunction in the Leigh syndrome mouse model with global *ndufs4* deletion.^32^ This may suggest that microglia are of less importance than infiltrating macrophages in driving the pathogenesis of neurodegeneration in mitochondrial diseases.

Finally, it is important to consider that pexidartinib and other drugs targeting CSF-1R do so by functioning as tyrosine kinase inhibitors, raising the possibility of off-target effects. Tyrosine kinases are involved in multiple cellular signaling pathways, including those related to the cell cycle, metabolism, and cell proliferation, differentiation, and survival.^40^ While potently inhibiting CSF-1R, pexidartinib is also known to inhibit the tyrosine kinases c-Kit (which likely accounts for its effect on hair color) and FLT3.^41,42^ It is clear that future experiments employing alternative genetic and pharmacological methods to achieve specific ablation of the various subpopulations of MNPs will be important to determine whether the salutary impact of pexidartinib in the *ndufs4* knockout model is truly a direct consequence of MNP depletion.

In summary, the combination of pexidartinib and hypoxia resulted in an additive, complete anatomic rescue of RGCs with complex I dysfunction. While the precise cellular mechanisms by which each treatment was neuroprotective in this model remain to be determined, the additive effect of the treatments demonstrates that there are at least two independent mechanisms leading to RGC survival. Our observations raise the intriguing prospect that a multimodal approach may be key to optimal management of mitochondrial optic neuropathies like LHON. Although reversing the mitochondrial impairment in RGCs that drives the optic neuropathy is a worthy therapeutic goal, modulation of neuroinflammation may be of similar importance in preventing optic atrophy.

## REFERENCES

1 Yu-Wai-Man P, Griffiths PG, Hudson G, Chinnery PF. Inherited Mitochondrial Optic Neuropathies. J Med Genet. 2009;46:145–158.

2 Sundaramurthy S, SelvaKumar A, Ching J, Dharani V, Sarangapani S, Yu-Wai-Man P. Leber Hereditary Optic Neuropathy-New Insights and Old Challenges. Graefes Arch Clin Exp Ophthalmol. 2021;259:2461–2472.

3 Borrelli E, Berni A, Cascavilla ML, et al. Visual Outcomes and Optical Coherence Tomography Biomarkers of Vision Improvement in Patients with Leber Hereditary Optic Neuropathy Treated with Idebenone. Am J Ophthalmol. 2023;247:35–41.

4 Klopstock T, Yu-Wai-Man P, Dimitriadis K, et al. A Randomized Placebo-Controlled Trial of Idebenone in Leber’s Hereditary Optic Neuropathy. Brain. 2011;134:2677–2686.

5 Yu-Wai-Man P, Carelli V, Newman NJ, et al. Therapeutic Benefit of Idebenone in Patients with Leber Hereditary Optic Neuropathy: The Leros Nonrandomized Controlled Trial. Cell Rep Med. 2024;5:101437.

6 Newman NJ, Biousse V, Yu-Wai-Man P, et al. Meta-Analysis of Treatment Outcomes for Patients with M.11778g>a Mt-Nd4 Leber Hereditary Optic Neuropathy. Surv Ophthalmol. 2025;70:283–295.

7 Newman NJ, Yu-Wai-Man P, Subramanian PS, et al. Randomized Trial of Bilateral Gene Therapy Injection for M.11778g>a Mt-Nd4 Leber Optic Neuropathy. Brain. 2023;146:1328–1341.

8 Wang L, Klingeborn M, Travis AM, Hao Y, Arshavsky VY, Gospe SM, 3rd. Progressive Optic Atrophy in a Retinal Ganglion Cell-Specific Mouse Model of Complex I Deficiency. Sci Rep. 2020;10:16326.

9 Lake NJ, Compton AG, Rahman S, Thorburn DR. Leigh Syndrome: One Disorder, More Than 75 Monogenic Causes. Ann Neurol. 2016;79:190–203.

10 Kruse SE, Watt WC, Marcinek DJ, Kapur RP, Schenkman KA, Palmiter RD. Mice with Mitochondrial Complex I Deficiency Develop a Fatal Encephalomyopathy. Cell Metab. 2008;7:312–320.

11 Quintana A, Kruse SE, Kapur RP, Sanz E, Palmiter RD. Complex I Deficiency Due to Loss of Ndufs4 in the Brain Results in Progressive Encephalopathy Resembling Leigh Syndrome. Proc Natl Acad Sci U S A. 2010;107:10996–11001.

12 Song L, Yu A, Murray K, Cortopassi G. Bipolar Cell Reduction Precedes Retinal Ganglion Neuron Loss in a Complex 1 Knockout Mouse Model. Brain Res. 2017;1657:232–244.

13 Warwick AM, Bomze HM, Wang L, et al. Continuous Hypoxia Reduces Retinal Ganglion Cell Degeneration in a Mouse Model of Mitochondrial Optic Neuropathy. Invest Ophthalmol Vis Sci. 2022;63:21.

14 Aguilar K, Comes G, Canal C, Quintana A, Sanz E, Hidalgo J. Microglial Response Promotes Neurodegeneration in the Ndufs4 Ko Mouse Model of Leigh Syndrome. Glia. 2022;70:2032–2044.

15 Stokes JC, Bornstein RL, James K, et al. Leukocytes Mediate Disease Pathogenesis in the Ndufs4(Ko) Mouse Model of Leigh Syndrome. JCI Insight. 2022;7.

16 Guo M, Schwartz TD, Dunaief JL, Cui QN. Myeloid Cells in Retinal and Brain Degeneration. FEBS J. 2022;289:2337–2361.

17 Margeta MA, Yin Z, Madore C, et al. Apolipoprotein E4 Impairs the Response of Neurodegenerative Retinal Microglia and Prevents Neuronal Loss in Glaucoma. Immunity. 2022;55:1627–1644 e1627.

18 Ritch MD, Hannon BG, Read AT, et al. Axonet: A Deep Learning-Based Tool to Count Retinal Ganglion Cell Axons. Sci Rep. 2020;10:8034.

19 Butowski N, Colman H, De Groot JF, et al. Orally Administered Colony Stimulating Factor 1 Receptor Inhibitor Plx3397 in Recurrent Glioblastoma: An Ivy Foundation Early Phase Clinical Trials Consortium Phase Ii Study. Neuro Oncol. 2016;18:557–564.

20 Wang Y, Zhao X, Gao M, et al. Myosin 1f-Mediated Activation of Microglia Contributes to the Photoreceptor Degeneration in a Mouse Model of Retinal Detachment. Cell Death Dis. 2021;12:926.

21 Berve K, West BL, Martini R, Groh J. Sex- and Region-Biased Depletion of Microglia/Macrophages Attenuates Cln1 Disease in Mice. J Neuroinflammation. 2020;17:323.

22 Shi Y, Manis M, Long J, et al. Microglia Drive Apoe-Dependent Neurodegeneration in a Tauopathy Mouse Model. J Exp Med. 2019;216:2546–2561.

23 Bolea I, Gella A, Sanz E, et al. Defined Neuronal Populations Drive Fatal Phenotype in a Mouse Model of Leigh Syndrome. Elife. 2019;8.

24 Jain IH, Zazzeron L, Goli R, et al. Hypoxia as a Therapy for Mitochondrial Disease. Science. 2016;352:54–61.

25 Benner B, Good L, Quiroga D, et al. Pexidartinib, a Novel Small Molecule Csf-1r Inhibitor in Use for Tenosynovial Giant Cell Tumor: A Systematic Review of Pre-Clinical and Clinical Development. Drug Des Devel Ther. 2020;14:1693–1704.

26 Chen BS, Newman NJ. Clinical Trials in Leber Hereditary Optic Neuropathy: Outcomes and Opportunities. Curr Opin Neurol. 2025;38:79–86.

27 Lyseng-Williamson KA. Idebenone: A Review in Leber’s Hereditary Optic Neuropathy. Drugs. 2016;76:805–813.

28 Li X, Yuan J, Qi J, et al. The Raav2-Nd1 Gene Therapy for Leber Hereditary Optic Neuropathy. Graefes Arch Clin Exp Ophthalmol. 2025.

29 Jain IH, Zazzeron L, Goldberger O, et al. Leigh Syndrome Mouse Model Can Be Rescued by Interventions That Normalize Brain Hyperoxia, but Not Hif Activation. Cell Metab. 2019;30:824–832 e823.

30 Meisel JD, Miranda M, Skinner OS, et al. Hypoxia and Intra-Complex Genetic Suppressors Rescue Complex I Mutants by a Shared Mechanism. Cell. 2024;187:659–675 e618.

31 Yu AK, Song L, Murray KD, et al. Mitochondrial Complex I Deficiency Leads to Inflammation and Retinal Ganglion Cell Death in the Ndufs4 Mouse. Hum Mol Genet. 2015;24:2848–2860.

32 Hanaford AR, Khanna A, Truong V, et al. Peripheral Macrophages Drive Cns Disease in the Ndufs4(-/-) Model of Leigh Syndrome. Brain Pathol. 2023;33:e13192.

33 Yu C, Roubeix C, Sennlaub F, Saban DR. Microglia Versus Monocytes: Distinct Roles in Degenerative Diseases of the Retina. Trends Neurosci. 2020;43:433–449.

34 O’Koren EG, Yu C, Klingeborn M, et al. Microglial Function Is Distinct in Different Anatomical Locations During Retinal Homeostasis and Degeneration. Immunity. 2019;50:723–737 e727.

35 Henry RJ, Ritzel RM, Barrett JP, et al. Microglial Depletion with Csf1r Inhibitor During Chronic Phase of Experimental Traumatic Brain Injury Reduces Neurodegeneration and Neurological Deficits. J Neurosci. 2020;40:2960–2974.

36 Qu W, Johnson A, Kim JH, Lukowicz A, Svedberg D, Cvetanovic M. Inhibition of Colony-Stimulating Factor 1 Receptor Early in Disease Ameliorates Motor Deficits in Sca1 Mice. J Neuroinflammation. 2017;14:107.

37 Wang W, Li Y, Ma F, et al. Microglial Repopulation Reverses Cognitive and Synaptic Deficits in an Alzheimer’s Disease Model by Restoring Bdnf Signaling. Brain Behav Immun. 2023;113:275–288.

38 Lei F, Cui N, Zhou C, Chodosh J, Vavvas DG, Paschalis EI. Csf1r Inhibition by a Small-Molecule Inhibitor Is Not Microglia Specific; Affecting Hematopoiesis and the Function of Macrophages. Proc Natl Acad Sci U S A. 2020;117:23336–23338.

39 O’Koren EG, Mathew R, Saban DR. Fate Mapping Reveals That Microglia and Recruited Monocyte-Derived Macrophages Are Definitively Distinguishable by Phenotype in the Retina. Sci Rep. 2016;6:20636.

40 Lemmon MA, Schlessinger J. Cell Signaling by Receptor Tyrosine Kinases. Cell. 2010;141:1117–1134.

41 Lamb YN. Pexidartinib: First Approval. Drugs. 2019;79:1805–1812.

42 Moss KG, Toner GC, Cherrington JM, Mendel DB, Laird AD. Hair Depigmentation Is a Biological Readout for Pharmacological Inhibition of Kit in Mice and Humans. J Pharmacol Exp Ther. 2003;307:476–480.

